# Epigenetic reprogramming by TET enzymes impacts co-transcriptional R-loops

**DOI:** 10.1101/2021.04.26.441414

**Authors:** João C. Sabino, Madalena R. de Almeida, Patricia L. Abreu, Ana M. Ferreira, Marco M. Domingues, Nuno C. Santos, Claus M. Azzalin, Ana R. Grosso, Sérgio F. de Almeida

## Abstract

DNA oxidation by ten-eleven translocation (TET) family enzymes is essential for epigenetic reprogramming. The conversion of 5-methylcytosine (5mC) into 5-hydroxymethylcytosine (5hmC) initiates developmental and cell-type-specific transcriptional programs through mechanisms that include changes in the chromatin structure. Here, we show that the presence of 5hmC in the transcribed DNA promotes the annealing of the nascent RNA to its template DNA strand, leading to the formation of an R-loop. The genome-wide distribution of 5hmC and R-loops show a positive correlation in mouse and human embryonic stem cells and overlap in half of all active genes. Moreover, R-loop resolution leads to differential expression of a subset of genes that are involved in crucial events during stem cell proliferation. Altogether, our data reveal that epigenetic reprogramming via TET activity promotes co-transcriptional R-loop formation, and disclose novel links between R-loops and the regulation of gene expression programs in stem cells.

## INTRODUCTION

During transcription, the nascent RNA molecule can hybridize with the template DNA and form a DNA:RNA hybrid and a displaced DNA strand. These triple-stranded structures, called R-loops, are physiologically relevant intermediates of several processes, such as immunoglobulin class-switch recombination and gene expression^1^. However, non-scheduled or persistent R-loops constitute an important source of DNA damage, namely DNA double-strand breaks (DSBs)^1^. To preserve genome integrity, cells possess diverse mechanisms to prevent the formation of R-loops or resolve them. R-loop formation is restricted by RNA-binding proteins and topoisomerase 1, whereas R-loops are removed by ribonucleases and helicases (reviewed in^1^). The ribonuclease H enzymes RNase H1 and RNase H2 degrade R-loops by digesting the RNA strand of the DNA:RNA hybrid. DNA and RNA helicases unwind the hybrid and restore the double-stranded DNA (dsDNA) structure. Several helicases unwind R-loops at different stages of the transcription cycle and in distinct physiological contexts^1^. For instance, we previously reported that the DEAD-box helicase 23 (DDX23) resolves R-loops formed during transcription elongation to regulate gene expression programs and prevent transcription-dependent DNA damage^2^. Intrinsic features of the transcribed DNA also influence its propensity to form R-loops. The presence of introns, for instance, prevents unscheduled R-loop formation at active genes^3^. An asymmetrical distribution of guanines (G) and cytosines (C) nucleotides in the DNA duplex also influences R-loop propensity, with an excess of Cs in the template DNA strand (positive G:C skew) favouring R-loop formation^4^. Moreover, chromatin and DNA features such as histone modifications, DNA-supercoiling and G-quadruplex structures also affect R-loop establishment^1^. R-loops can also drive chromatin modifications. Promoter-proximal R-loops enhance the recruitment of the Tip60–p400 histone acetyltransferase complex and inhibit the binding of polycomb repressive complex 2 and histone H3 lysine-27 methylation^5^. R-loops formed over G-rich terminator elements promote histone H3 lysine-9 dimethylation, a repressive mark that reinforces RNA polymerase II pausing during transcription termination^6–8^.

Besides affecting histone modifications, R-loops also act as barriers against DNA methylation spreading into active genes^4,9^. DNA methylation, namely 5-methylcytosine (5mC), results from the covalent addition of a methyl group to the carbon 5 of a C attached to a G through a phosphodiester bond (CpG)^10^. The activity of DNA methyltransferase (DNMT) enzymes makes 5mC widespread across the mammalian genome where it plays major roles in imprinting, suppression of retrotransposon silencing and gene expression^11^. More than 70% of all human gene promoters contain stretches of CpG dinucleotides, termed CpG islands (CGIs), whose transcriptional activity is repressed by CpG methylation^11,12^. R-loops positioned near the promoters of active genes maintain CGIs in an unmethylated state^9^, likely by reducing the affinity of DNMT1 binding to DNA^13^, or recruiting methylcytosine dioxygenases ten-eleven translocation (TET) enzymes^14^.

The TET enzymes family members share the ability to oxidize 5mC to 5-hydroxymethylcytosine (5hmC)^15,16^. 5hmC is a relatively rare DNA modification found across the genome much less frequently than 5mC^17^. Genome-wide, 5hmC is more abundant at regulatory regions near transcription start sites (TSSs), promoters and exons, consistent with its role in gene expression regulation^18^. The levels of 5hmC are enriched at active promoter regions, as observed upon activation of neuronal function-related genes in neural progenitors and neurons^15,19^. 5hmC has the potential to modify the DNA helix structure by favouring DNA-end breathing motion, a dynamic feature of the protein–DNA complexes thought to control DNA accessibility^17^. Moreover, 5hmC weakens the interaction between DNA and nucleosomal H2A-H2B dimers, facilitating RNA polymerase II elongation, and diminishes the thermodynamic stability of the DNA duplex^17^. While 5mC increases the melting temperature, 5hmC reduces the amount of energy needed to separate the two strands of the DNA duplex^20,21^. Molecular dynamics simulations revealed that the highest amplitude of GC DNA base-pair fluctuations is observed in the presence of 5hmC, whereas 5mC yielded GC base pairs with the lower amplitude values^21^. The presence of 5hmC destabilizes GC pairing by alleviating steric constraints through an increase in molecular polarity^21^.

Because features that destabilize the DNA duplex, such as supercoiling or G-quadruplexes, are known to facilitate nascent RNA annealing with the template DNA strand, we reasoned that 5hmC may favour R-loop formation. Here, we show that 5hmC promotes R-loop formation during *in vitro* transcription of DNA templates. Moreover, changing the expression levels and genomic targeting of TET enzymes affects R-loop levels in cells. Analysis of genome-wide distribution profiles shows a positive correlation between 5hmC and R-loops in mouse embryonic stem (mES) and human embryonic kidney 293 (HEK293) cells, with a clear overlap of 5hmC and R-loops in approximately half of all active genes. We also show that 5hmC-rich regions are characterized by increased levels of phosphorylated histone H2AX (γH2AX), a marker of DNA damage. Finally, by determining the pathways more significantly affected by R-loops formed at 5hmC loci, we propose a novel function for R-loops in regulating gene expression programs that drive stem cell proliferation.

## RESULTS

### Transcription through 5hmC-rich DNA favours R-loop formation

To assess the impact of cytosine methylation on R-loop formation, we performed *in vitro* T7 transcription of DNA fragments containing either native or modified cytosine deoxyribonucleotides (dCTPs). We synthesized three distinct DNA transcription templates, each composed of a T7 promoter followed by a 400bp sequence containing a genomic region prone to form R-loops *in vivo*^2,8^. Two of these sequences (*β-actin* P1 and *β-actin* P2) are from the transcription termination region of the *β-actin* gene; the third sequence is from the *APOE* gene. The DNA templates for the *in vitro* transcription reactions were generated by PCR-amplification in the presence of dNTPs containing either native C, 5mC, or 5hmC (**Figure 1A**). Successful incorporation of dCTP variants was confirmed by immunoblotting using specific antibodies against each variant (**Figure 1B**). The formation of R-loops during the *in vitro* transcription reactions was inspected by blotting immobilized RNAs with the S9.6 antibody, which recognizes the DNA:RNA hybrids (**Figure 1C**). To increase the specificity of hybrid detection, all samples were treated with RNase A in high salt conditions in order to digest all RNA molecules except those engaged in R-loops. The specific detection of DNA:RNA hybrids was confirmed by blotting transcription reaction products previously digested with RNase H (**Figure 1C**). In agreement with our hypothesis that 5hmC favours R-loops, increased amounts of DNA:RNA hybrids were detected in samples derived from *in vitro* transcription of 5hmC-rich *β-actin* P1, *β-actin* P2 and *APOE* DNA templates (**Figure 1D)**. To directly visualize and quantify R-loop structures obtained in the *in vitro* transcription reactions we performed atomic force microscopy (AFM) experiments (**Figure 1E**). R-loops were visualized as blob, spur or loop structures, as previously described^22,23^. Quantification of these structures revealed that transcription products from 5hmC-rich DNA templates were enriched in R-loops, which were extensively lost upon RNase H treatment.

**Figure 1:**
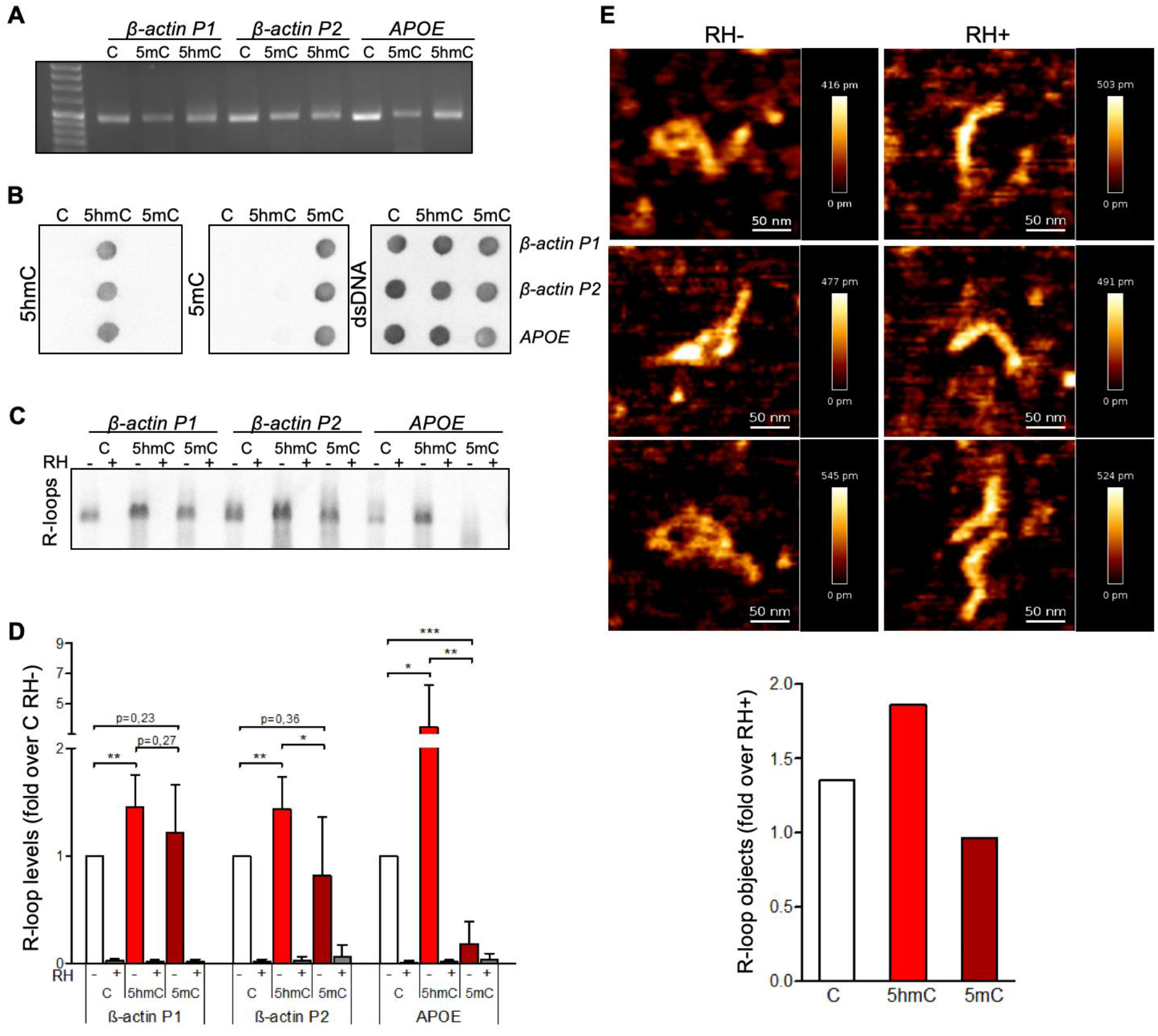
5hmC favours co-transcriptional R-loop formation. **(A)** Native or modified dCTPs were incorporated upon PCR amplification into DNA fragments with sequences from the transcription termination region of the β-*actin* gene (β-*actin* P1 and β-*actin* P2) or the *APOE* gene. **(B)** Incorporation of dCTP variants confirmed by immunoblotting using specific antibodies against 5mC, 5hmC and double-stranded DNA (dsDNA). **(C)** R-loops formed upon *in vitro* transcription reactions were detected by immunoblotting using the S9.6 antibody. RNase H-treated *in vitro* transcription reaction products (RH+) serve as negative controls. All data are representative of seven independent experiments with similar results. **(D)** S9.6 immunoblots were quantified and the R-loop levels normalized against the levels detected in the reaction products of DNA templates containing native C. Data represent the mean and standard deviation (SD) from seven independent experiments. *p<0.05, **p<0.01 and ***p<0.001, using two-tailed Student’s t test. **(E)** *In vitro* transcription reaction products of β*-actin* P2 templates were visualized using atomic force microscopy. R-loop structures obtained from 5hmC-containing β*-actin* P2 transcription in the absence (RH-) or presence (RH+) of RNase H are shown. R-loops present in the transcription reaction products of C, 5mC or 5hmC-containing β-*actin* P2 templates were counted in a minimum of 80 filaments observed in three individual AFM experiments.

### TET enzymatic activity impacts endogenous R-loop levels

We next sought to test whether the 5hmC DNA modification induces R-loop formation in cells. We quantified R-loop levels in mES cells carrying doxycycline (dox)-inducible shRNAs targeting either *Tet1* or *Tet3*^24^ (**Figure 2A**). In agreement with their role in converting 5mC into 5hmC, knockdown of *Tet1* and *Tet3* in mES cells resulted in decreased total cellular 5hmC, whereas 5mC showed a mild increase (**Figure 2B**). Dot-blot hybridization of total cellular nucleic acids using the S9.6 antibody revealed that depletion of each TET enzyme reduced endogenous R-loops (**Figure 2C**). We then asked whether global changes in transcription rates contributed to reducing R-loop levels in TET1-depleted cells. We quantified fluorescent 5-ethynyl uridine (EU) incorporation into nascent RNA molecules in mES cells upon TET1 knockdown (**Figure 2D**). TET1 knockdown did not reduce global EU incorporation, indicating that diminished transcription cannot account for the observed reduction in R-loop levels.

**Figure 2:**
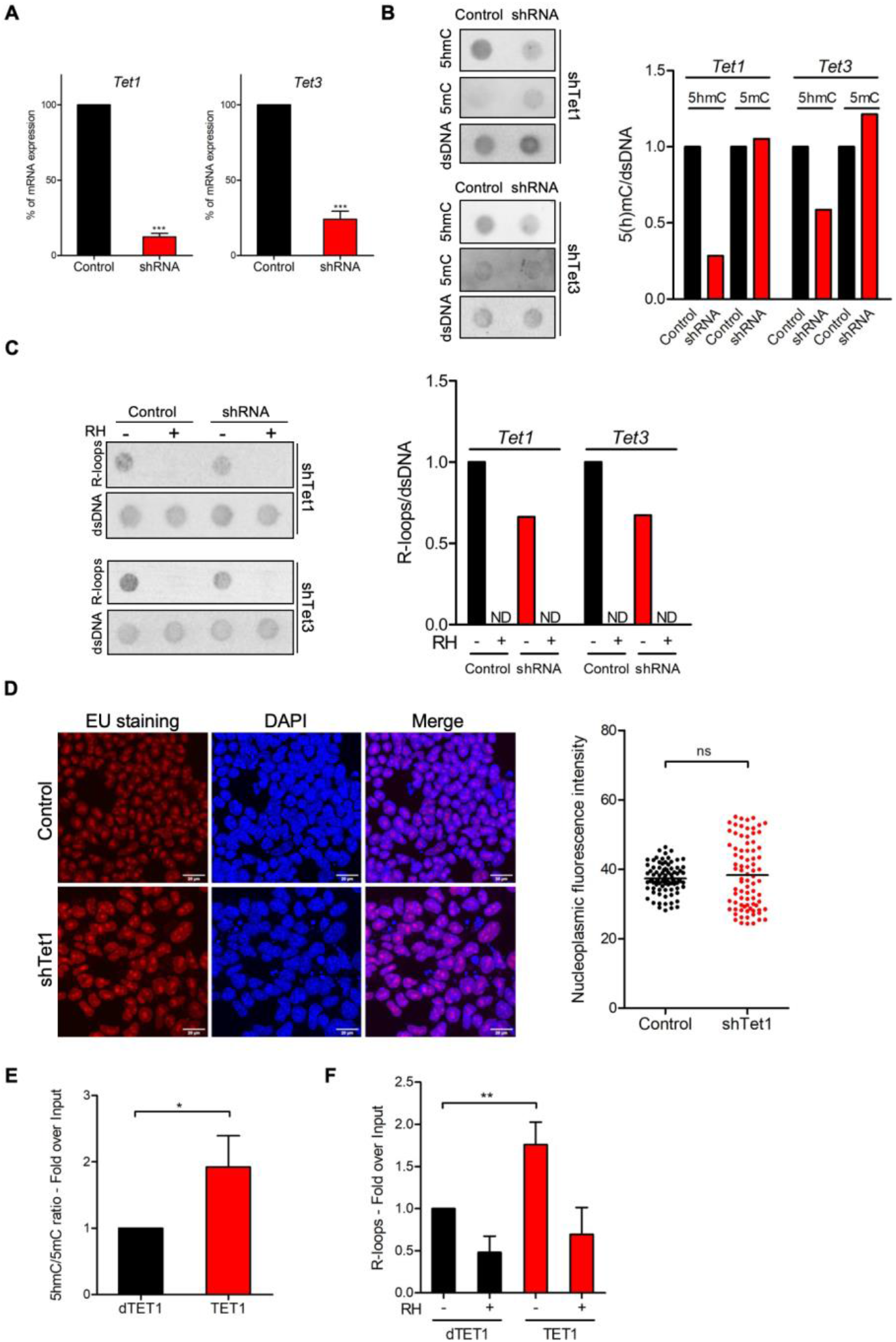
TET enzymatic activity impacts R-loop levels. **(A)** *Tet1* and *Tet3* mRNA expression levels in mES cells stably expressing dox-inducible shRNA targeting *Tet1* or *Tet3*. Graphs show mean and SD of mRNA expression in dox-treated cells normalized to control cells (no dox). Data are from three independent experiments. ***p<0.001, two-tailed Student’s t test. **(B)** 5mC and 5hmC immunoblot in *Tet1* and *Tet3*-depleted cell extracts (shTet1 and shTet3). Blots were quantified, normalized against dsDNA levels and plotted in a bar graph. Data are representative of three independent experiments. **(C)** R-loops were detected by immunoblot in *Tet1* and *Tet3*-depleted cell extracts. Blots were quantified, normalized against dsDNA levels and plotted in a bar graph. Data are representative of three independent experiments. ND=not detected. **(D)** Transcription levels in *Tet1*-depleted mES cells assessed through EU-incorporation. DAPI was added to stain DNA. Scale bars: 20 μm. Data are representative of three independent experiments with similar results. The scatter plot represents EU nucleoplasmic fluorescence intensity. Horizontal solid lines represent the mean values. Over 90 cells from three individual experiments were scored for each experimental condition and statistical significance was determined using the Mann Whitney test (p>0.05). **(E)** Changes in the 5hmC/5mC ratio at the last exon of the *APOE* gene in U2OS cells expressing dCas9-TET1 or dCas9-dTET1 with gRNA targeting this locus. Data show the mean and SD of the 5hmC/5mC ratio obtained in cells expressing dCas9-TET1 normalized to the same ratio in dCas9-dTET1 cells from three independent experiments. * p<0.05, two-tailed Student’s t test. **(F)** R-loop levels in the APOE locus assessed by DRIP. Data show R-loop levels in dCas9-TET1 expressing cells normalized to dCas9-dTET1 cells. RNase H-treated samples (RH+) serve as negative controls. Mean and SD are from three independent DRIP experiments. **p<0.01, two-tailed Student’s t test.

Next, we employed a modified CRISPR-based system to target TET enzymatic activity to specific loci^25^. We used a pool of three specific guide RNAs (gRNAs) to direct a catalytically inactive Cas9 nuclease fused to the catalytic domain of TET1 (dCas9-TET1) to the last exon of the *APOE* gene. As a control, dCas9 was fused to an inactive mutant version of the TET1 catalytic domain (dCas9-dTET1). Local enrichment of 5hmC following dCas9-TET1 targeting at the *APOE* locus was confirmed by DNA immunoprecipitation using antibodies specific for 5mC or 5hmC modified nucleotides (**Figures 2E**). DNA:RNA immunoprecipitation (DRIP) experiments detected increased R-loop levels in the last exon of *APOE* upon tethering of dCas9-TET1 but not of dCas9-dTET1 (**Figures 2F**). Collectively, these data suggest that editing 5hmC density by changing the expression levels or the genomic distribution of TET enzymes influences R-loop formation in cells.

### 5hmC and R-loops overlap genome-wide at transcriptionally active genes

To further inspect the link between 5hmC and R-loops we performed computational analyses of 5hmC antibody-based DNA immunoprecipitation (hMeDIP-seq) and DNA:RNA immunoprecipitation (DRIP-seq) datasets from mES and HEK293 cells^5,26–28^. To assess individual genome-wide distribution profiles, R-loops density was probed over fixed windows of ±10 kbp around the 5hmC peaks (**Figure 3A and Supplementary Figure 1A**). The resulting metagene plots and heatmaps revealed a marked overlap between 5hmC-rich loci and R-loops. Despite the distinct distribution patterns of 5hmC (well-defined peaks) and R-loops (reads spanning genomic regions with highly heterogeneous lengths, ranging between a few dozen to over 1 kb^5^, we could obtain a statistically significant Pearson correlation coefficient between both (p<0.05) (**Figure 3B and Supplementary Figure 1B**). Furthermore, approximately half of all R-loops detected genome-wide occurred at 5hmC-containing loci (**Figure 3C and Supplementary Figure 1C**). Notably, we observed an overlap between 5hmC and R-loops in 6839 (51%) out of the 13288 actively expressed genes (**Figure 3D**), a feature illustrated in the individual profiles of mouse and human genes (**Figure 3E and Supplementary Figure 1D**). Metagene profiles revealed very similar patterns of intragenic distribution, with both 5hmC and R-loops increasing towards the transcription termination site (TTS), where they reached maximum levels (**Figure 3F**). At the transcription start sites (TSS), however, the 5hmC DNA modification was mostly absent, whereas R-loops were abundant. The detection of R-loop peaks at TSS regions is in agreement with previous studies^4,9^ and imply that 5hmC is not necessary for co-transcriptional DNA:RNA hybridization and R-loop formation.

**Figure 3:**
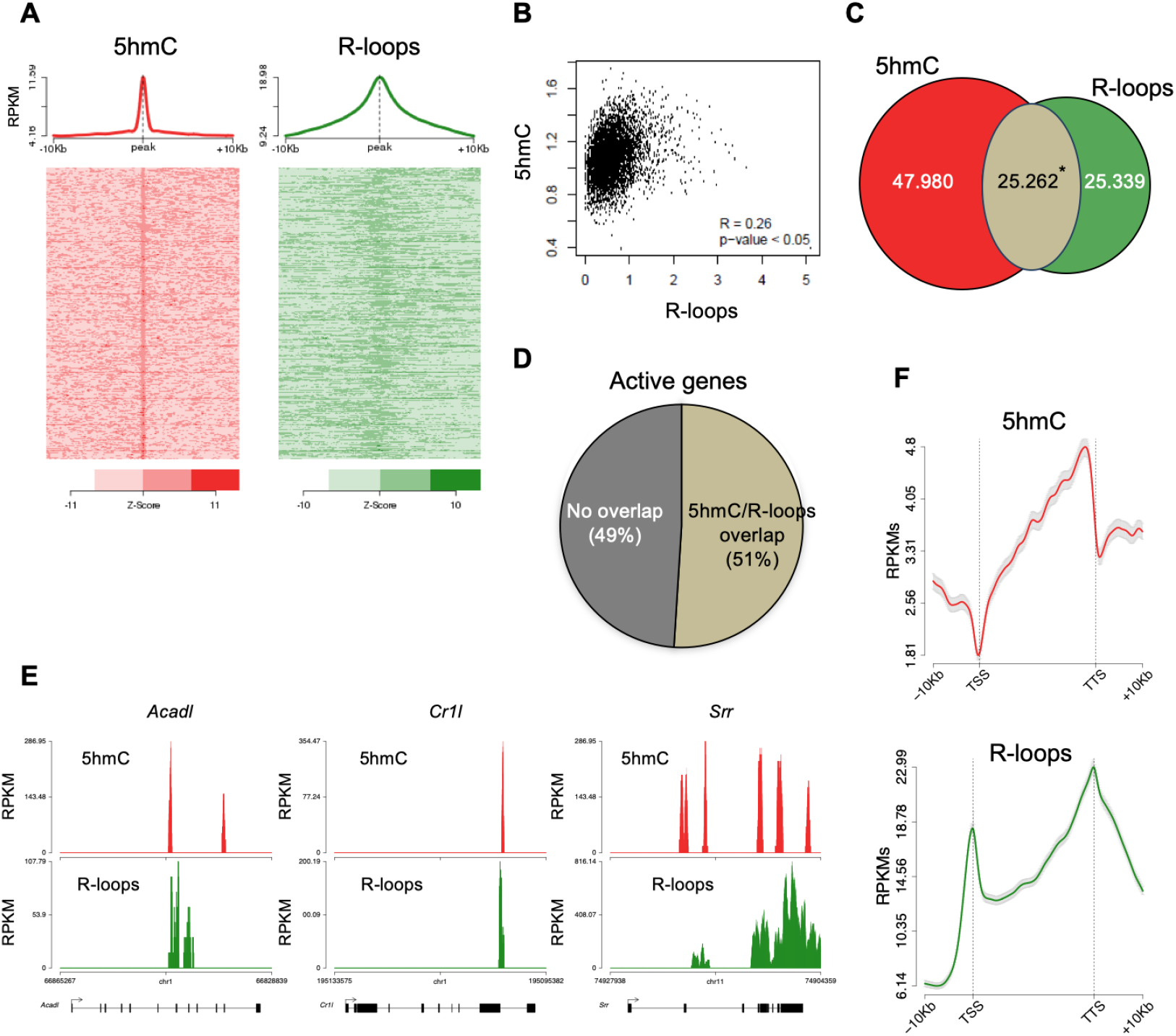
5hmC and R-loops overlap in active genes of mES cells. **(A)** Metagene and heatmap profiles of 5hmC and R-loops probed over fixed windows of ±10 kbp around the 5hmC peaks in expressed genes. **(B)** Pearson correlation coefficient between 5hmC and R-loops distribution within active genes (p<0.05). **(C)** Number of loci displaying 5hmC, R-loops, and overlapping 5hmC and R-loops. * Permutation analysis, p<0.05. **(D)** Percentage of active genes displaying overlapping 5hmC and R-loops. **(E)** Individual profiles of 5hmC and R-loop distribution along the *Acadl, Cr1l* and *Srr* genes. Density signals are represented as reads per kilobase (RPKMs). **(F)** Metagene profiles of 5hmC and R-loops distribution in active genes. The gene body region was scaled to 60 equally-sized bins and ±10 kbp gene-flanking regions were averaged in 200bp windows. TSS: transcription start site. TTS: transcription termination site. Density signals are represented as RPKMs and error bars (gray) represent standard error of the mean.

We then sought to simultaneously detect 5hmC and R-loops at the same loci in individual mES cells. We performed proximity ligation assays (PLA) using S9.6 and anti-5hmC antibodies (**Figure 4A**). While control reactions with each antibody alone or without primary antibodies did not produce a significant PLA signal, staining with S9.6 and anti-5hmC antibodies gave rise to a robust signal scattered throughout the nucleus in 92% of all cells (**Figure 4B**).

**Figure 4:**
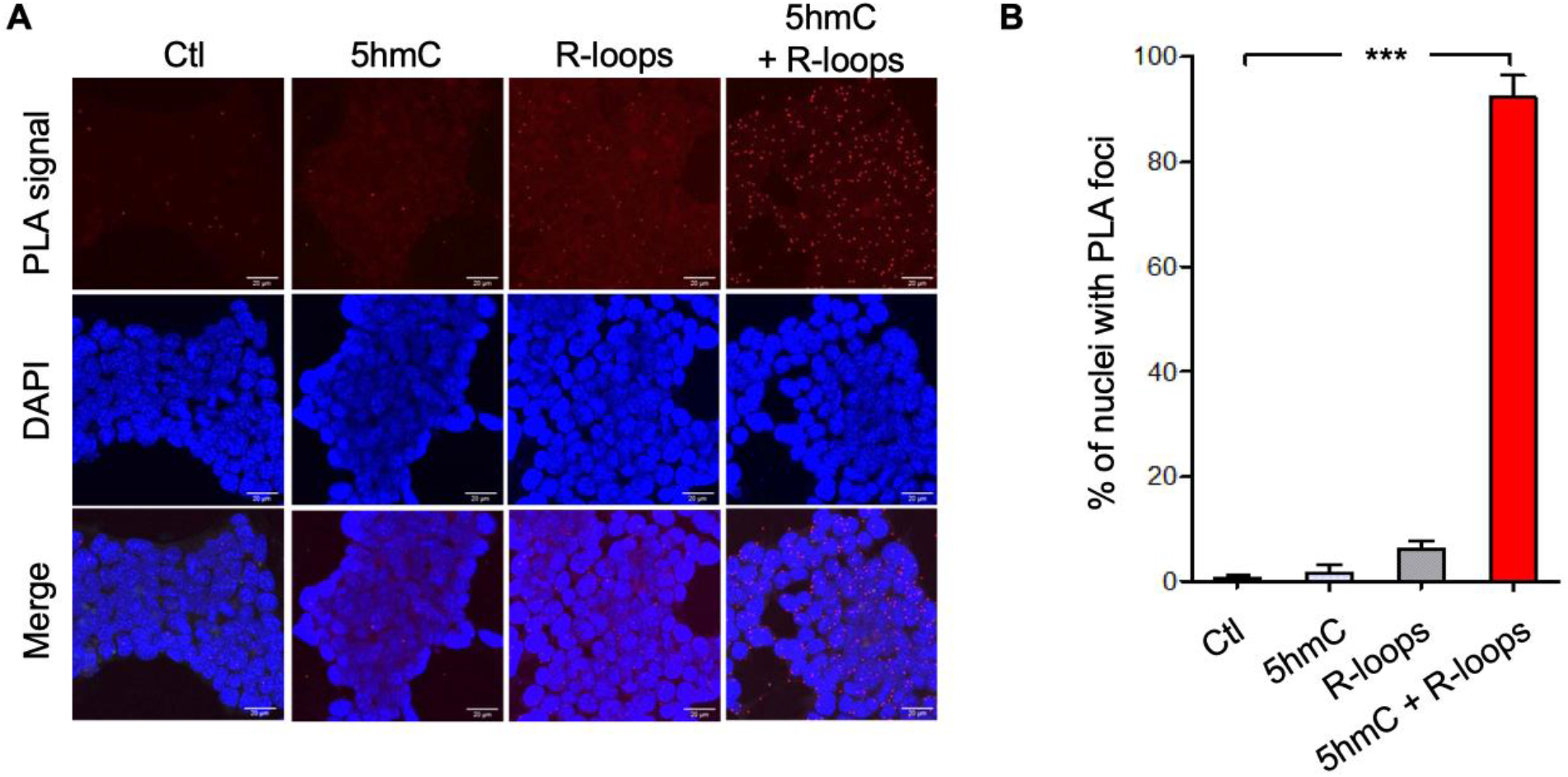
Simultaneous detection of 5hmC and R-loops at the same genomic loci in individual mES cells. **(A)** 5hmC and R-loops PLA foci in mES cells. DAPI was added to the mounting medium to stain DNA. Scale bars: 20μm. Data are representative of three independent experiments with similar results. **(B)** The bar graph shows the mean and SD of the percentage of cells containing 5hmC and R-loops PLA foci. A minimum of 300 cells from three individual experiments was scored for each experimental condition. ***p<0.001, two-tailed Student’s t test.

### 5hmC-rich loci are genomic hotspots for DNA damage

Disruption of R-loop homeostasis is a well-described source of genomic instability^1^. For instance, co-transcriptional R-loops increase conflicts between transcription and replication machineries by creating an additional barrier to fork progression^29,30^. Such conflicts may cause DNA damage, including DSBs, which can be revealed using antibodies against γH2AX. We analysed the genomic distribution of γH2AX by interrogating chromatin immunoprecipitation followed by sequencing (ChIP-seq) data from HEK293 cells^31^. The individual distribution profiles of γH2AX were analysed over fixed windows of ±10 kbp around the 5hmC peaks detected in the same cells (**Figure 5A**). The resulting metagene plots revealed marked enrichment of γH2AX at 5hmC-rich loci. The genic distribution of 5hmC and R-loops along three different genes further showed co-localization of the two marks with γH2AX (**Figure 5B**). Analysis of γH2AX and 5hmC distribution within active genes revealed a low yet statistically significant Pearson correlation coefficient (p<0.05) (**Figure 5C**).

**Figure 5:**
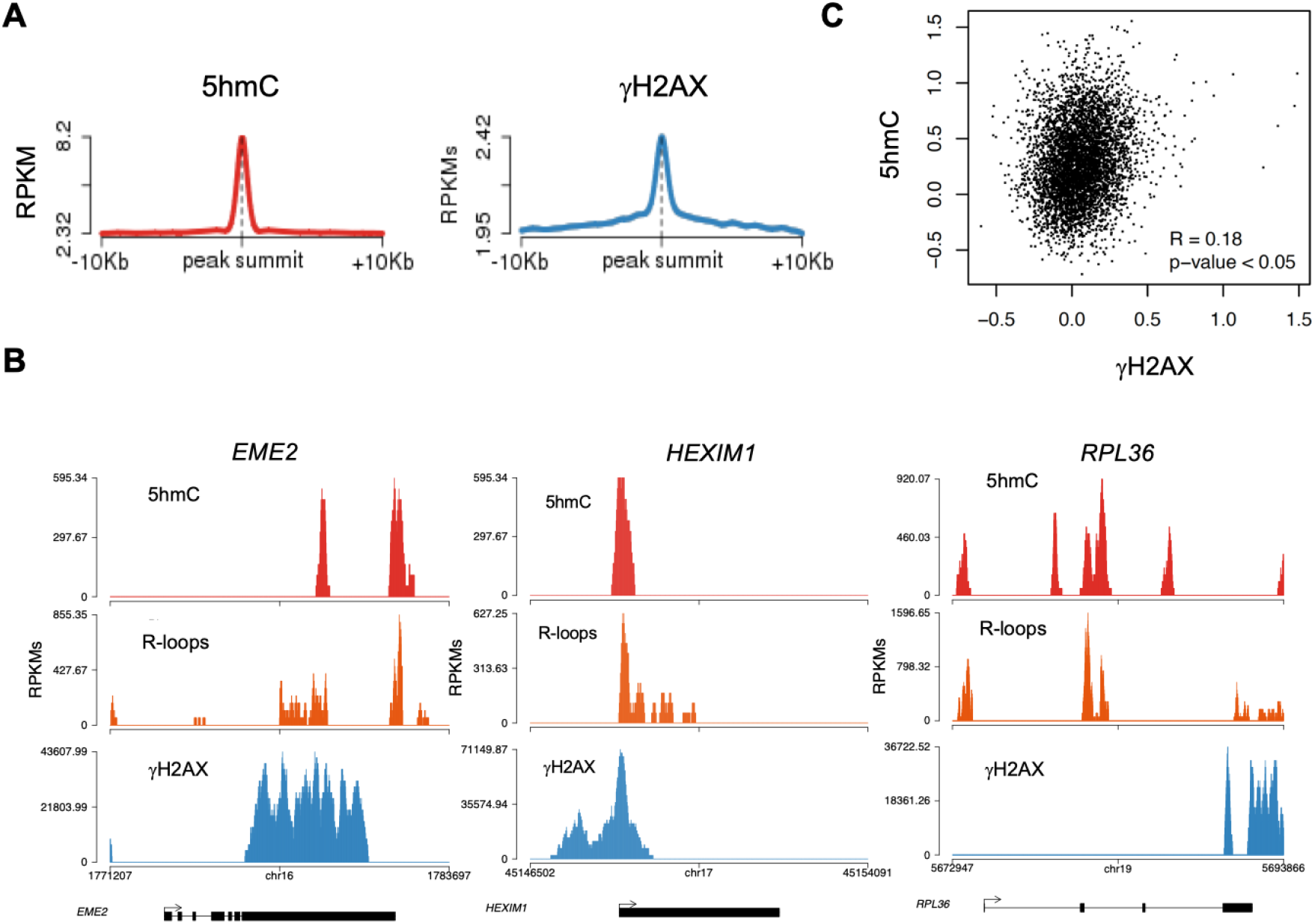
5hmC-rich loci are genomic hotspots for DNA damage. **(A)** Metagene profiles of 5hmC and γ H2AX probed over fixed windows of ±10 kbp around the 5hmC peaks in expressed genes of HEK293 cells. **(B)** Individual profiles of 5hmC, R-loops and γH2AX distribution along the *EME2, HEXIM1* and *RPL36* genes. Density signals are represented as reads per kilobase (RPKMs). **(C)** Pearson correlation coefficient between 5hmC and γH2AX at active genes (p<0.05).

### R-loops formed at 5hmC-rich regions impact the expression of genes involved in establishing diapause

To gather insights into the functional impact of R-loops at 5hmC-rich DNA regions, we analysed whole-transcriptome (RNA-seq) of mES cells overexpressing RNase H, a condition resulting in genome-wide loss of R-loops^5^. Amongst the genes that were differentially expressed, we found that 64% and 48% of all downregulated and upregulated genes, respectively, displayed R-loops overlapping with 5hmC (**Figure 6A**). Pathway analysis revealed that these differentially expressed genes (**Supplementary Table 1**) are involved in the mechanistic target of rapamycin (mTOR) (downregulated) and MYC (upregulated) signalling pathways (**Figure 6B and C)**. mTOR and MYC are known to play opposite roles in establishing diapause, the temporary suspension of embryonic development driven by adverse environmental conditions^32^, a stage that ES cells mimic when cultured *in vitro*. mTOR, a major nutrient sensor, acts as a rheostat during ES cell differentiation and reductions in mTOR activity trigger diapause^33^. This raises the hypothesis that RNase H impacts the proliferation of ES cells. To directly investigate this hypothesis, we overexpressed RNase H in mES cells. Analysis of the cellular DNA content 24 and 48h after RNase H overexpression did not reveal any significant changes in the cell cycle progression (**Supplementary Figure 2A and B**). This finding suggests that fine-tuned R-loop formation at specific loci, rather than global changes in R-loop levels, commands the activation of specific gene expression programs in ES cells.

**Figure 6:**
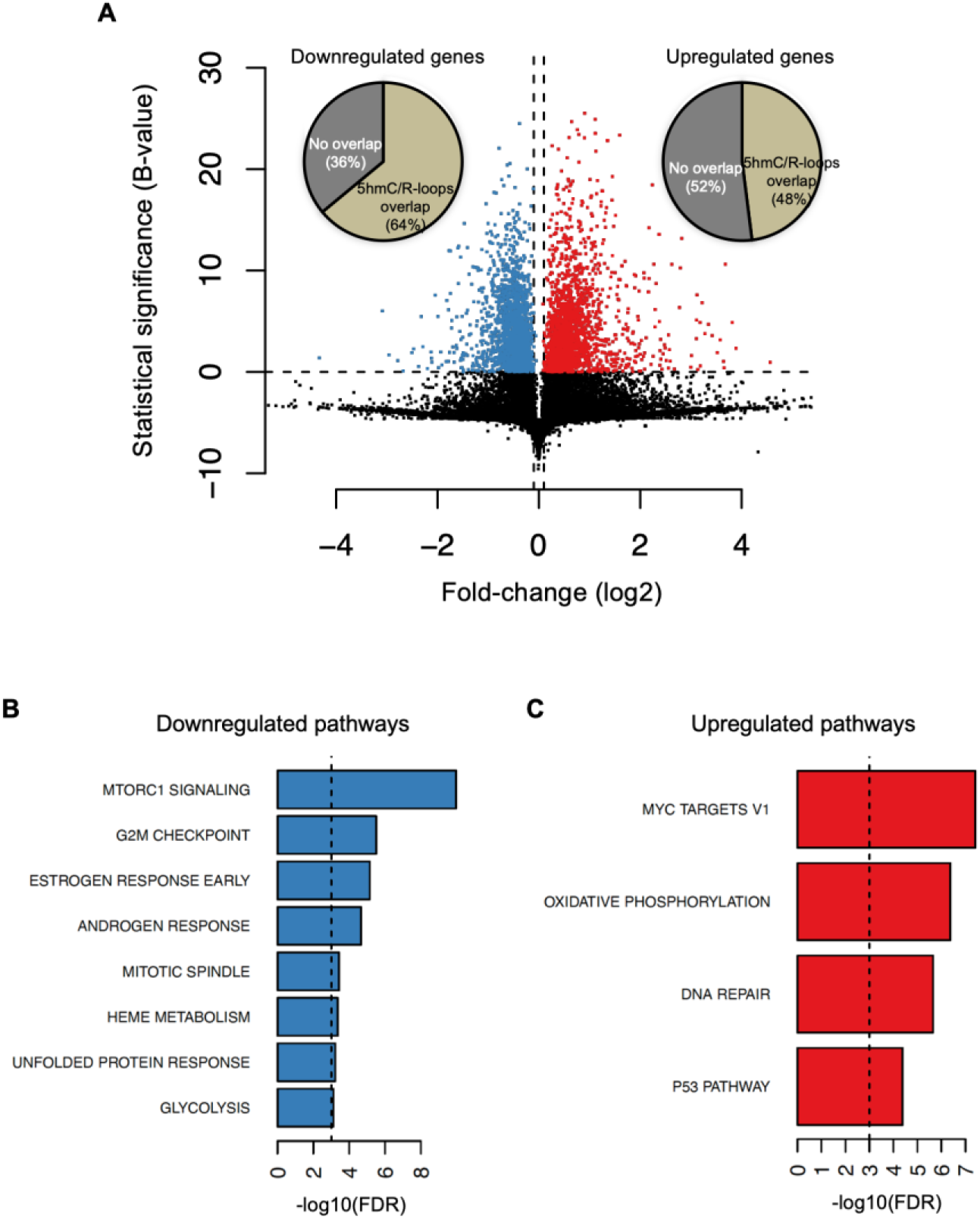
Cellular pathways affected by R-loops formed at 5hmC loci. **(A)** Volcano plot displaying the differentially expressed genes in mES cells upon RNase H overexpression. Of all downregulated and upregulated genes, 64% and 48% displayed R-loops overlapping with 5hmC, respectively. **(B-C)** Pathway analysis of the genes that have R-loops overlapping with 5hmC and are differentially expressed upon RNase H overexpression. Shown are the significantly downregulated **(B)** and upregulated **(C)** hallmark gene sets from MSigDB. False discovery rate (FDR), p<0.001.

## DISCUSSION

In this study, we probed the hypothesis that 5hmC facilitates the co-transcriptional formation of non-canonical DNA secondary structures, known as R-loops. Data from *in vitro* transcription reactions and atomic force microscopy provide direct evidence showing that transcription through 5hmC-rich DNA favours R-loop formation. Using a well-established cellular model that allows the selective depletion of TET enzymes from mES cells, we demonstrate that TET activity increases endogenous R-loop levels. Notably, the diminished levels of R-loops observed in TET-depleted cells did not result from impaired transcription, suggesting that 5hmC directly promotes R-loop formation. In agreement, tethering TET enzymes to a specific genomic locus using a CRISPR/Cas9-based system increase the levels of R-loops at the target locus.

As R-loops play diverse physiological roles^1^, our findings associate TET activity to numerous novel functions such as the regulation of gene expression, telomere homeostasis or the maintenance of genome integrity. Whether 5hmC-editing at promoter regions or gene 3’ ends instructs transcription initiation or termination, respectively, through the regulation of R-loops is still to be directly investigated. Nevertheless, our finding that genome-wide 5hmC and R-loops overlap more robustly at the transcription termination site of active genes supports a model whereby TET enzymes act upstream of R-loop formation during transcription termination^34^. Telomeres, the nucleoprotein complexes found at the ends of linear eukaryotic chromosomes, can be maintained in proliferating ES and cancer cells by either the activity of telomerase or the alternative lengthening of telomeres (ALT) pathway^35^. ALT telomeres are maintained by mechanisms relying on homologous recombination (HR) between telomeric repeats. R-loops form extensively during transcription of telomeric-repeat-containing RNA (TERRA) and trigger a telomere specific replication stress, which promotes HR and re-elongation of telomeres by ALT^36,37^. Notably, mES cells depleted of *Tet1* and/or *Tet2* exhibit short telomeres and chromosomal instability, concomitant with reduced telomere recombination^38^. This suggests that telomeric 5hmC might promote HR at telomeres through the establishment of R-loops.

Owing to their link with R-loops, 5hmC may also harm genome integrity if not properly controlled. Indeed, we found that 5hmC-rich loci are hotspots for DNA damage genome-wide. In addition to altering the expression levels of tumour suppressors or oncogenes^39^, our findings suggest that TET-driven changes in the DNA methylation landscape may as well drive transcription-dependent genome damaging events that could facilitate cancer development and progression. In agreement with this view, a TET1 isoform that lacks regulatory domains, including its DNA binding domain, but retains its catalytic activity is enriched in cancer cells^40^, suggesting that mis-targeted TET activity may drive oncogenic events, such as genomic instability. Conversely, TET activity deposits 5hmC at DNA damage sites induced by aphidicolin or microirradiation in HeLa cells and prevents chromosome segregation defects in response to replication stress^41^. Hence, TETs may play dual roles as both oncogenic and tumour suppressor genes, with the former arising as the consequence of altered expression levels or function, as observed in several cancers, such as triple-negative breast cancer^39,42^.

While the role that TET enzymes play during carcinogenesis is not yet clear, the impact of 5hmC on stem cell differentiation and development has been extensively studied^24^. By driving the developmental DNA methylome reprogramming, TETs carry out numerous functions related to early developmental processes. Here, we disclose a putative new role for R-loops as mediators of 5hmC-driven gene expression programs that determine the self-renewal and differentiation capabilities of stem cells. Our gene ontology analysis revealed that R-loops formed at 5hmC-rich regions impact the expression of genes involved in establishing diapause. This stage of temporary suspension of embryonic development is triggered by adverse environmental conditions^32^. Accordingly, changes in the activity of mTOR, a major nutrient sensor, control ES cell commitment to trigger diapause^33^. The mTOR signalling pathway was significantly downregulated upon global R-loop suppression by RNase H. Conversely, MYC targets, which prevent ES cells from entering the state of dormancy that characterizes diapause^43^, were amongst the genes more significantly upregulated upon RNase H overexpression in mES cells. MYC proteins drive hypertranscription in ES cells, accelerating the gene expression output associated with increased cell proliferation^44^. In agreement with the view that 5hmC-driven R-loop formation impacts functions related with mES cell proliferation, we observed a significant upregulation of genes involved in oxidative phosphorylation (OXPHOS), DNA repair and p53 signalling upon RNase H overexpression. Upregulation of OXPHOS, the main source of energy in most mammalian cells, including ES cells, may fulfil the energetic needs of ES cells resuming proliferation as they exit diapause^45^. Augmented expression of DNA repair and p53 signalling strengthen the genome caretaker and gatekeeper mechanisms that cope with the DNA damage burst observed in highly proliferative cells^46^. This seemingly dichotomous effect of RNase H overexpression in ES cells (i.e. decreased mTOR and increased MYC signalling, simultaneously), was further corroborated by the lack of significant changes in the proliferation rate of mES cells upon RNase H overexpression. Whether the controlled 5hmC-driven formation of R-loops at specific genes, namely MYC or mTOR targets, is sufficient to commit ES cells towards proliferation or establishing diapause and how do TET enzymes capture the environmental cues to target R-loop formation at selected genes are important questions that emerge from our findings. Thus, our data set the ground for further research aimed at investigating the role of R-loops in ES cells.

## MATERIALS AND METHODS

### Cell lines and culture conditions

E14TG2a (E14) mouse embryonic stem (mES) cells were provided by Domingos Henrique (Instituto de Medicina Molecular João Lobo Antunes), and were a gift from Austin Smith (Univ. of Exeter, UK)^47^. 129S4/SvJae (J1) mES cells were kindly provided by Joana Marques (Medical School, University of Porto). Cells were grown as monolayers on 0,1% gelatine (410875000, Acros Organics) coated dishes, using Glasgow Modified Eagle’s Medium (GMEM) (21710-025, Gibco), supplemented with 1% (v/v) 200mM L-glutamine (25030-024, Thermo Scientific), 1% (v/v) 100mM sodium pyruvate (11360-039, Gibco), 1% (v/v) 100x non-essential aminoacids solution (11140-035, Gibco), 0,1% (v/v) 0,1M 2-mercaptoethanol (M7522, Sigma Aldrich), 1% (v/v) penicillin-streptomycin solution (15070-063, Gibco) and 10% (v/v) heat-inactivated, ES-qualified FBS (SH30070, Cytiva). Medium was filtered through a 0,22μm filter. Home-produced leukaemia inhibitory factor (LIF) was added to the medium upon plating, at 6×10^−2^ ng/μL. U2OS osteosarcoma and HEK293T embryonic kidney cells (purchased from ATCC) were grown as monolayers in Dulbecco’s Modified Eagle medium (DMEM) (21969-035, Gibco), supplemented with 1% (v/v) 200mM L-glutamine (25030-024, Thermo Scientific), 1% (v/v) penicillin-streptomycin solution (15070-063, Gibco) and 10% (v/v) FBS (10270106, Gibco). All cells were maintained at 37°C in a humidified atmosphere with 5% CO_2_.

### *Tet* knockdown

J1 mES cells with doxycycline-inducible short hairpin RNA-micro RNA (shRNA – **Supplementary Table 2**) sequences targeting *Tet1* or *Tet3* were treated for 48h with 2 μg/mL doxycycline (D9891, Sigma Aldrich).

### RNA isolation and quantitative RT-PCR

Total RNA was isolated from J1 mES cells under doxycycline treatment for 48h, using TRIzol reagent (15596018, Invitrogen). cDNA was prepared through reverse transcriptase activity (MB125, NZYTech). RT-qPCR was performed in the ViiA 7 Real-Time PCR system (Applied Biosystems), using PowerUp SYBR Green Master Mix (A25918, Applied Biosystems). The relative RNA expression was estimated as follows: 2^(Ct reference - Ct sample)^, where Ct reference and Ct sample are mean threshold cycles of RT-qPCR done in duplicate for the *U6* snRNA or *Gapdh* and for the gene of interest, respectively. Primer sequences are presented in **Supplementary Table 3**.

### R-loops dot blot

J1 mES cells were collected after 48h of doxycycline treatment and lysed in lysis buffer (100mM NaCl, 10mM Tris pH 8.0, 25mM EDTA pH 8.0, 0,5% SDS, 50 μg/mL Proteinase K) overnight at 37°C. Nucleic acids were extracted using standard phenol-chloroform extraction protocol and re-suspended in DNase/RNase-free water. Nucleic acids were then fragmented using a restriction enzyme cocktail (20U each of EcoRI, BamHI, HindIII, BsrgI and XhoI). Half of the sample was digested with 40U RNase H (MB085, NZYTech) to serve as negative control, for about 36-48h at 37ºC. Digested nucleic acids were cleaned with standard phenol-chloroform extraction and res-suspended in DNase/RNase-free water. Nucleic acids samples were quantified in a NanoDrop 2000 spectrophotometer (Thermo Scientific), and equal amounts of DNA were deposited into a positively charged nylon membrane (RPN203B, GE Healthcare). Membranes were UV-crosslinked using UV Stratalinker 2400 (Stratagene), blocked in 5% (m/v) milk in PBSt (PBS 1× containing 0.05% (v/v) Tween 20) for 1h at room temperature, and immunoblotted with specific antibodies. Details of antibodies used are included in **Supplementary Table 4**.

### 5mC and 5hmC dot blot

J1 mES cells were collected after 48h of doxycycline treatment and lysed in lysis buffer (100mM NaCl, 10mM Tris pH 8.0, 25mM EDTA pH 8.0, 0.5% SDS, 50 μg/mL Proteinase K) for 2h at 56°C. Samples were sonicated with 4-6 pulses of 15s at 10mA intensity using a Soniprep150 sonicator (keeping tubes for at least 1min on ice between pulses) to shear chromatin into 100-300bp fragments. Fragmented nucleic acids were cleaned with standard phenol-chloroform extraction method and re-suspended in DNase/RNase-free water. DNA was resolved in agarose gels to confirm fragment size. Samples were denatured by boiling at 100°C for 10min, followed by immediate chilling on ice and quick spin, and deposited into a nylon membrane (the sample fraction used for dsDNA detection was not subject to boiling), prior to UV-crosslinking and immunoblotting. Details of antibodies used are included in **Supplementary Table 4**.

### 5-ethynyl uridine (EU) staining

J1 mES cells were grown on glass coverslips and incubated for 1h (37°C, 5% CO_2_) with EU from the Click-iT RNA Alexa Fluor 594 imaging kit (C10330, Invitrogen). Cells were fixed with 3,7% formaldehyde in PBS 1× for 15min at room temperature and permeabilized with 0.5% Triton X-100 in PBS 1× for 15min at room temperature. The Click-It reaction using a fluorescent azide (Alexa Fluor 594 azide) was then performed according to manufacturer’s instructions (30min at room temperature, protected from light). Finally, nuclear staining was performed with Hoechst 33342 1:1000 in PBS 1× for 10min at room temperature, and coverslips were assembled in Vectashield (Vector Laboratories) mounting medium. Cells were imaged using a point-scanning confocal microscope Zeiss LSM 880, 63×/1.4 oil immersion, with stacking acquisition and generation of maximum intensity projection images. Nucleoplasmic fluorescence intensity measurements were performed using ImageJ.

### Proximity Ligation Assay (PLA)

E14 mES cells were grown on coverslips for 48h, and fixed/permeabilized with methanol for 10min on ice, followed by 1min acetone on ice. Cells were then incubated with both primary antibodies simultaneously for 1h at 37ºC, followed by a pre-mixed solution of PLA probe anti-mouse minus (DUO92004, Sigma Aldrich) and PLA probe anti-rabbit plus (DUO92002, Sigma Aldrich) for 1h at 37° C. Localized rolling circle amplification was performed using Detection Reagents Red (DUO92008, Sigma Aldrich), according to the manufacturer’s instructions. Slides were mounted in 1:1000 DAPI in Vectashield. Images were acquired using the Point Scanning Confocal Microscope Zeiss LSM 880, 63x/1,4 oil immersion, with stacking acquisition and generation of maximum intensity projection images. The number of PLA foci was quantified using ImageJ. Details of antibodies used are mentioned in **Supplementary Table 4**.

### g-blocks PCR

Designed g-blocks were ordered from IDT (**Supplementary Table 5**), and PCR-amplified using Phusion High-Fidelity DNA Polymerase (M0530S, NEB), according to manufacturer’s instructions. M13 primers were used to amplify all fragments (**Supplementary Table 3**), in the presence of dNTP mixes containing native (MB08701, NZYTech), methylated (D1030, Zymo Research) or hydroxymethylated (D1040, Zymo Research) cytosines. Efficient incorporation of modified dCTPs was confirmed through immunoblotting with specific antibodies. Details of antibodies used are mentioned in **Supplementary Table 4**.

### *In vitro* transcription

PCR products were subject to *in vitro* transcription using the HiScribe T7 High Yield RNA Synthesis Kit (E2040S, NEB), which relies on the T7 RNA polymerase to initiate transcription from a T7 promoter sequence (present in our fragments). Reactions were performed for 2h at 37°C, using 1 μg of DNA as template, according to manufacturer’s instructions.

### S9.6 immunoblotting of *in vitro* transcription products

Half of each *in vitro* transcription product was treated with 10U RNase H (MB085, NZYTech) at 37°C overnight, to serve as negative control. Then, all samples were treated with 0,05U RNase A (10109142001, Roche) at 350mM salt concentration, for 15min at 37ºC, and ran on agarose gel. Nucleic acids were transferred to a nylon membrane through capillary transfer, overnight at room temperature. The membrane was then UV-crosslinked twice, blocked in 5% milk in PBSt for 1h at room temperature, and incubated with the primary antibody at 4ºC overnight. Signal quantification was performed using Image Lab. Details of antibodies used are included in **Supplementary Table 4**.

### Atomic Force Microscopy

RNase A-treated *in vitro* transcription products, treated or not with RNase H, were purified through phenol-chloroform extraction method and re-suspended in DNase/RNase-free water. DNA solution was diluted 1:10 in Sigma ultrapure water (with final 10mM MgCl_2_) and briefly mixed to ensure even dispersal in solution. A 10μL droplet was deposited at the centre of a freshly cleaved mica disc, ensuring that the pipette tip did not contact the mica substrate. The solution was let to adsorb on mica surface for 1-2min to ensure adequate coverage. The mica surface was carefully rinsed with Sigma ultrapure water, so that excess of poorly bound DNA to mica is removed from the mica substrate. Afterwards, the mica substrate was dried under a gentle stream of argon gas for approximately 2min, making sure that any excess water is removed. DNA imaging was performed using a JPK Nanowizard IV atomic force microscope, mounted on a Zeiss Axiovert 200 inverted optical microscope. Measurements were carried out in tapping mode using commercially available ACT cantilevers (AppNano). After selecting a region of interest, the DNA was scanned in air, with scan rates between 0.5 and 0.9 Hz. The setpoint selected was close to 0.3 V. Several images from different areas of the same sample were performed and at least three independent samples for each condition were imaged. All images were of 512 × 512 pixels and analysed with JPK data processing software.

### Lentiviral transduction

Lentivirus containing dCas9-TET1 (#84475, Addgene) or dCas9-dTET1 (#84479, Addgene) coding plasmids, as well as one out of three gRNAs (gRNA_1, 2 and 3) coding plasmids designed for the *APOE* last exon, were produced. HEK293T cells were transfected with the above-mentioned plasmids, as well as with the Δ8.9 and VSV-g plasmids (for virus assembly). Virus production occurred for 48h, after which culture supernatant was collected and filtered through a 0.45μm filter. Lentivirus were collected through ultracentrifugation (25000 rpm, 3h, 4°C) using a SW-41Ti rotor in a Beckman XL-90 ultracentrifuge. Virus were re-suspended in PBS 1× and stored at −80°C. For infection, a pool of lentivirus containing dCas9-TET1 or dCas9-dTET1, as well as gRNA_1, 2 or 3 coding plasmids, was used to infect seeded U2OS cells. After 24h, antibiotic selection was performed with 1.5 μg/mL puromycin, and infection proceeded for more 48h. 3 days post-infection, cells were harvested and genomic DNA was extracted for subsequent protocols.

### DNA:RNA Immunoprecipitation (DRIP)

Infected U2OS cells were collected and lysed in lysis buffer (100mM NaCl, 10mM Tris pH 8.0, 25mM EDTA, 0.5% SDS, 50μg/mL Proteinase K) overnight at 37°C. Nucleic acids were extracted using standard phenol-chloroform extraction protocol and re-suspended in DNase/RNase-free water. Nucleic acids were then fragmented using a restriction enzyme cocktail (20U each of EcoRI, BamHI, HindIII, BsrgI and XhoI), and 10% of the digested sample was kept aside to use later as input. Half of the remaining volume was digested with 40U RNase H (MB085, NZYTech) to serve as negative control, for 72h at 37°C. Digested nucleic acids were cleaned with standard phenol-chloroform extraction and re-suspended in DNase/RNase-free water. RNA:DNA hybrids were immunoprecipitated from total nucleic acids using 5µg of S9.6 antibody (MABE1095, Merck Millipore) in binding buffer (10mM Na_2_HPO_4_ pH 7.0, 140mM NaCl, 0.05% Triton X-100), overnight at 4°C. 50µl protein G magnetic beads (10004D, Invitrogen) were used to pull-down the immune complexes at 4°C for 2-3h. Isolated complexes were washed 5 times (for 1 min on ice) with binding buffer and once with Tris-EDTA (TE) buffer (10mM Tris pH 8.1, 1mM EDTA). Elution was performed in two steps, for 15min at 55°C each, using elution buffer (50mM Tris pH 8.0, 10mM EDTA, 0.5% SDS, 60µg/mL Proteinase K). The relative occupancy of DNA:RNA hybrids was estimated by RT-qPCR as follows: 2^(Ct Input−Ct^ ^IP)^, where Ct Input and Ct IP are mean threshold cycles of RT-qPCR done in duplicate for input samples and specific immunoprecipitations, respectively. Primer sequences are presented in **Supplementary Table 3**.

### 5-(hydroxy)Methylated DNA Immunoprecipitation ((h)MeDIP)

Infected U2OS cells were collected and lysed in lysis buffer overnight at 37°C. Samples were sonicated with 4 pulses of 15s at 10mA intensity using a Soniprep150 sonicator (keeping tubes for at least 1min on ice between pulses). Fragmented nucleic acids were cleaned with standard phenol-chloroform extraction protocol and res-suspended in DNase/RNase-free water. 10% of sample was kept aside to use later as input. The remaining volume was denatured by boiling the samples at 100°C for 10min, followed by immediate chilling on ice and quick spin. Samples were divided in half, and 5µg of anti-5mC antibody (61255, Active Motif) or 5µg of anti-5hmC antibody (39791, Active Motif) were used to immunoprecipitate 5mC and 5hmC, respectively, in binding buffer, overnight at 4°C. 50µl protein G magnetic beads (10004D, Invitrogen) were used to pull-down the immune complexes at 4°C for 2-3h. Isolated complexes were washed 5 times (for 1 min on ice) with binding buffer and once with TE buffer. Elution was performed in two steps, for 15min at 55°C each, using elution buffer. The relative occupancy of 5mC and 5hmC was estimated by RT-qPCR. Primer sequences are presented in **Supplementary Table 3**.

### Cell cycle analysis

pEGFP-N1 (GFP coding plasmid used as control) was purchased from Addgene, and pEGFP-RNaseH1 (GFP-tagged RNase H1 coding plasmid) was kindly provided by Robert J. Crouch (NIH, USA). Seeded mES cells were transfected with GFP (control) or GFP-tagged RNase H coding plasmids. 24 or 48h later, cells were trypsinized and pelleted by centrifugation at 500×*g* for 5min. Cells were fixed in cold 1% PFA for 20min at 4°C, followed by permeabilization in 70% ethanol for 1h at 4°C. Cells were then treated with 25 μg/mL RNase A (10109142001, Roche) in PBS 1× at 37 °C for 20min, followed by staining with 20 μg/mL propidium iodide (P4864, Sigma Aldrich) in PBS 1× for 10 min at 4°C. Flow cytometry was performed on a BD Accuri C6 (BD Biosciences) and data were analysed using FlowJo software.

### Multi-omics data

High-throughput sequencing (HTS) data for mES cells and HEK293 cells were gathered from GEO archive: transcriptome of mES cells (GSE67583); R-loops in mES cells (GSE67581); 5hmC in mES cells (GSE31343); γ H2AX in mES cells (GSE69140); active transcription in HEK293 (GRO-seq, GSE51633); R-loops in HEK293 (DRIP-seq, GSE68948); 5hmC modification in HEK293 (hMeDIP-seq, GSE44036); γ H2AX (ChIP-seq, GSE75170). Transcriptome profiles of mES cells overexpressing RNase H were obtained from GSE67583. The quality of HTS data was assessed with FastQC (www.bioinformatics.babraham.ac.uk/projects/fastqc).

### 5hmC, R-loop and γ H2AX genome-wide characterization

The HTS datasets produced by immunoprecipitation (DRIP-seq, ChIP-seq and hMeDIP-seq) were analysed through the same workflow. First, the reads were aligned to the reference mouse and human genome (mm10 and GRCh38/hg38 assemblies, respectively) with Bowtie^48^, and filtering for uniquely aligned reads. Enriched regions were identified relative to the input samples using MACS^49^, with a false-discovery rate of 0,05. Finally, enriched regions were assigned to annotated genes, including a 4-kilobase region upstream the transcription start site and downstream the transcription termination site. Gene annotations were obtained from mouse and human Gencode annotations (M11 and v23 versions, respectively) and merged into a single transcript model per gene using BedTools^50^. For individual and metaprofiles, uniquely mapped reads were extended in the 3’ direction to reach 150 nt with the Pyicos^51^. Individual profiles were produced using a 20bp window. For the metaprofiles centered around 5hmC peaks: 5hmc enriched regions were aligned by the peak summit (maximum of the peak) and the read density for the flanking 10 kbp were averaged in a 200bp window. For the metagene profiles: the gene body region was scaled to 60 equally sized bins and ±10 kbp gene-flanking regions were averaged in 200bp windows. All profiles were plotted as normalized reads per kilobase per million mapped reads (RPKMs). A set of in-house scripts for data processing and graphical visualization were written in bash and in the R environmental language http://www.R-project.org^52^. SAMtools^53^ and BEDtools were used for alignment manipulation, filtering steps, file format conversion and comparison of genomic features. Statistical significance of the overlap between 5hmC regions and R-loops was assessed by permutation analysis. Briefly, random 5hmC and R-loops datasets were generated 1000 times from annotated genes using the shuffle BEDtools function (maintaining the number and length of the originally datasets). The p-value was determined as the frequency of overlapping regions between the random datasets as extreme as the observed.

### Transcriptome analysis

Expression levels (Transcripts per Million, TPMs) from RNA-seq and GRO-seq datasets were obtained using Kallisto^54^, where reads were pseudo-aligned to mouse and human Gencode transcriptomes (M11 and v23, respectively). Transcriptionally active genes for 5hmC and R-loops annotation were defined as those with expression levels higher than the 25^th^ percentile. Differential expression in mES cells overexpressing RNase H was assessed using edgeR (v3.20.9) and limma (v3.34.9) R packages^55,56^. Briefly, samples comparison was performed using voom transformed values, linear modelling and moderated T-test as implemented in limma R package, selecting significantly differentially expressed genes with B-statistics higher than zero. Significantly enriched pathways of up and down-regulated genes (with overlapping R-loops/5hmC regions) were selected using Fisher’s Exact Test and all expressed genes as background gene list. Evaluated pathways were obtained from the hallmark gene sets of Molecular Signatures Database (MSigDB)^57^ and filtered using False discovery rate corrected p-values < 0,05.

## ACKNOWLEDGMENTS

We thank our colleagues, Joana Marques, Domingos Henrique and Robert Crouch for kind gifts of cell lines, plasmids and reagents. This work was funded by PTDC/BIA-MOL/30438/2017 and PTDC/MED-OUT/4301/2020 from Fundação para a Ciência e Tecnologia (FCT), Portugal. Funding was also received from EU Horizon 2020 Research and Innovation Programme (RiboMed 857119). J.C.S. is the recipient of an FCT PhD fellowship PD/BD/128292/2017.

---

**Supplementary Figure 1:**
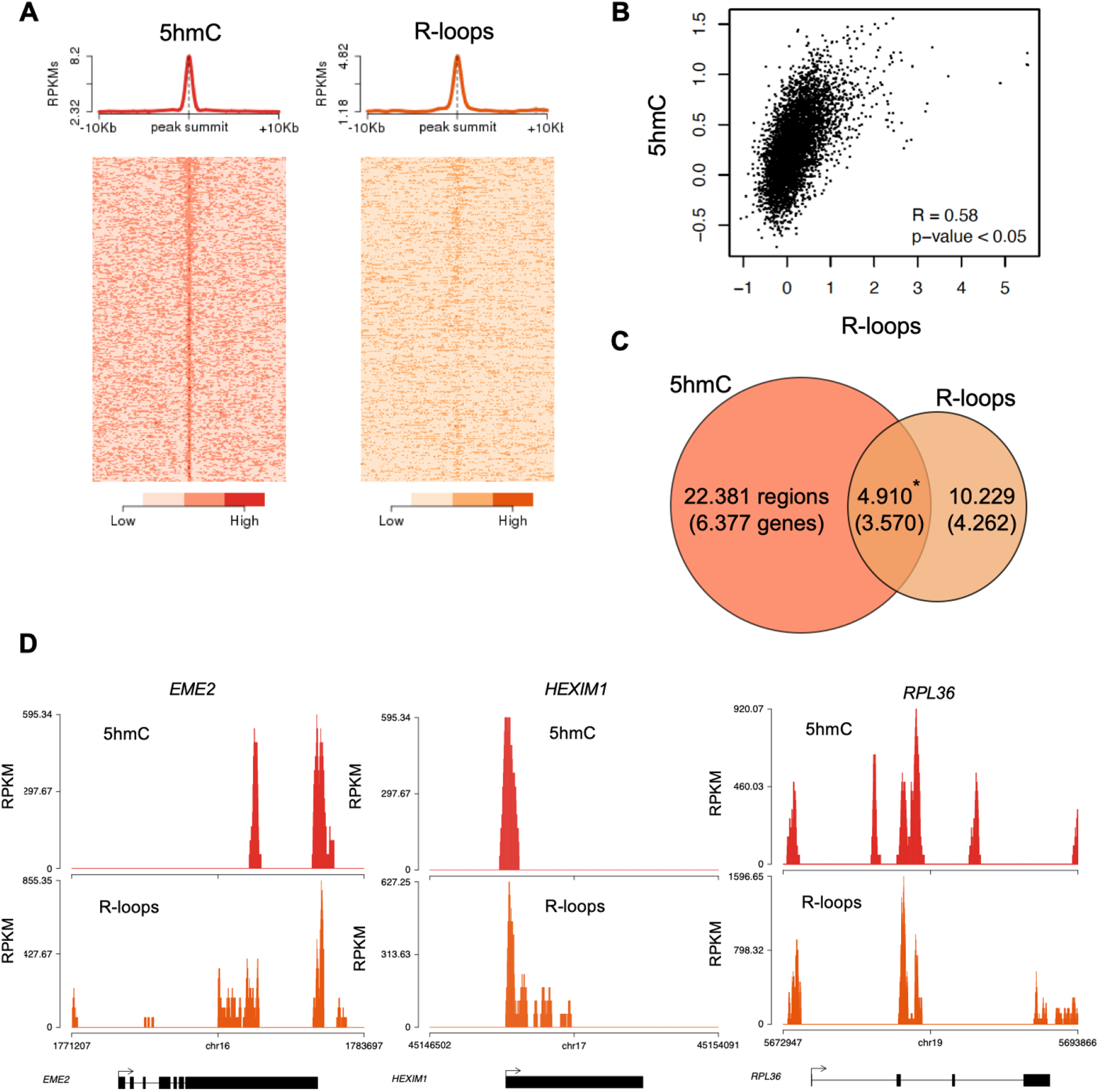
Genome-wide analysis of 5hmC and R-loops in HEK293 cells. **(A)** Metagene and heatmap profiles of 5hmC and R-loops probed over fixed windows ±10 kbp around the 5hmC peaks in expressed genes. **(B)** Pearson correlation coefficient between 5hmC and R-loops distribution within active genes (p<0.05). **(C)** Number of loci displaying 5hmC, R-loops, and overlapping 5hmC and R-loops. * Permutation analysis, p<0.05. **(D)** Individual profiles of 5hmC and R-loop distribution along the *EME2, HEXIM1* and *RPL36* genes. Density signals are represented as reads per kilobase (RPKMs).

**Supplementary Figure 2:**
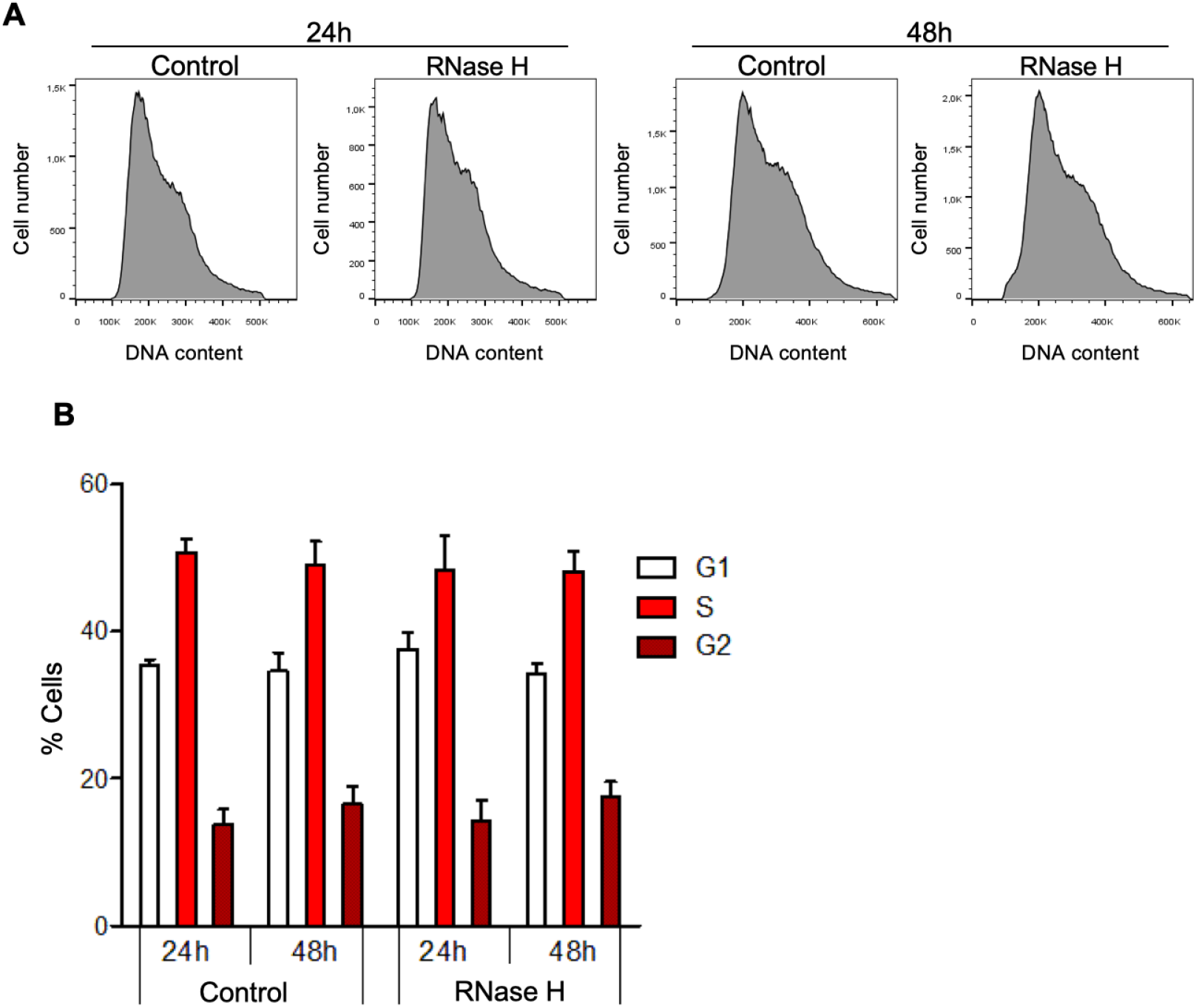
Global R-loop suppression does not impact cell cycle progression of mES cells. **(A)** Flow cytometry analysis of propidium iodide-treated mES cells with ectopic expression of either GFP (control) or GFP-tagged RNase H for 24 or 48h. Data are representative of five independent experiments. **(B)** Percentage of control and RNase H-overexpressing mES cells at each cell cycle stage. Means and SDs are from five independent experiments.

## Source data figure legends

**Figure 1 – source data 1**. Original, uncropped images of all blots shown in Figure 1.

**Figure 2 – source data 1**. Original, uncropped images of all blots shown in Figure 2.

**Supplementary Table 1: Differentially expressed genes upon RNase H overexpression**. (attached Excel file)

**Supplementary Table 2:**
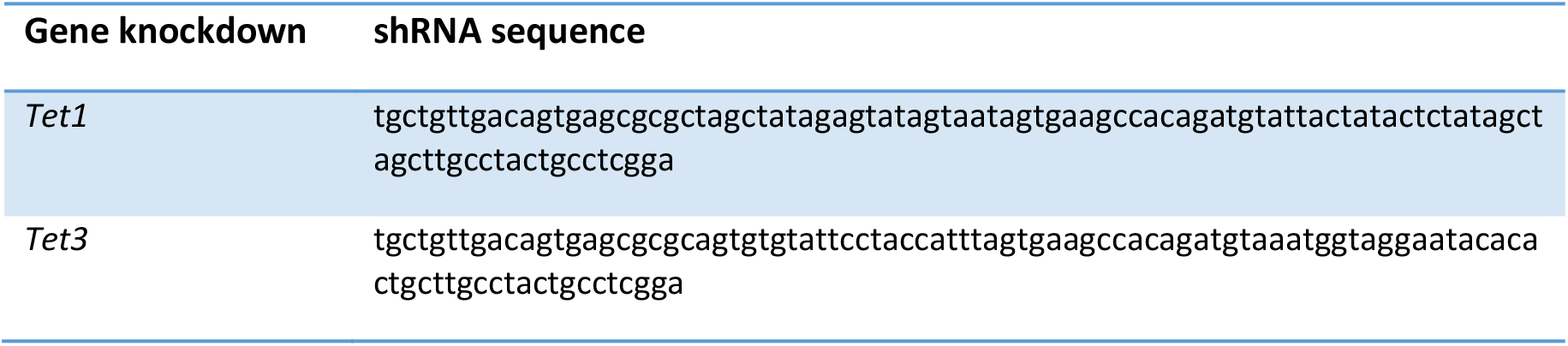
shRNA sequences.

**Supplementary Table 3:**
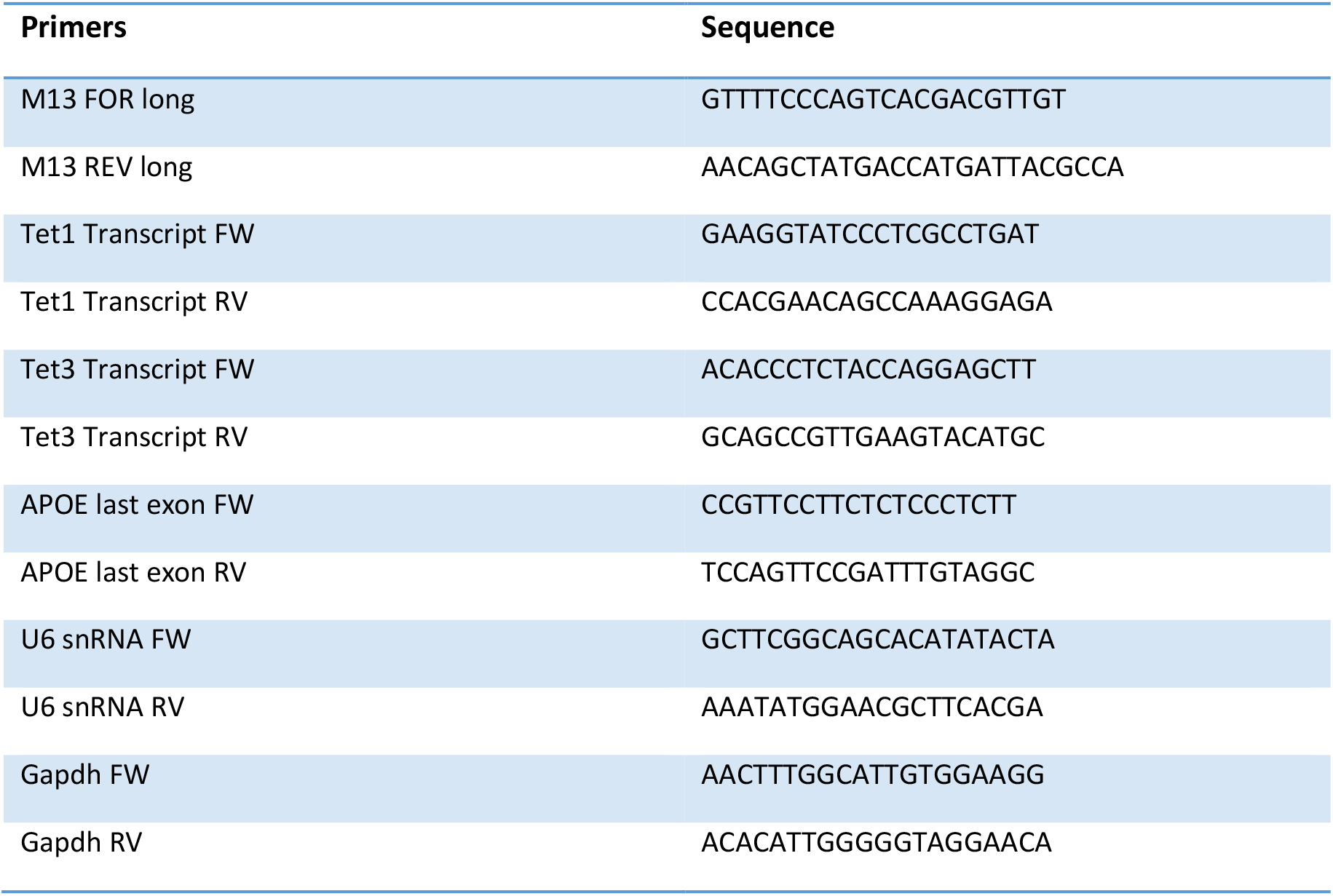
Oligonucleotide sequences.

**Supplementary Table 4:** Antibodies used in this study.

| Product | Concentrations | Company/ Cat. No. | Notes |
| --- | --- | --- | --- |
| S9.6 | 5ug/ IP; 1:1000 (DB) | Millipore; MABE1095 | Anti-DNA:RNA hybrid antibody used to detect R-loops |
| dsDNA | 1:1000 (DB) | Santa Cruz; sc-58749 | Anti-dsDNA specific antibody (HYB331-01) |
| 5hmC | 5ug/ IP; 1:1000 (DB) | Active Motif; 39791 | 5-hydroxymethylcytosine antibody |
| 5mC | 5ug/ IP; 1:500 (DB) | Active Motif; 61255 | 5-methylcytosine antibody |

**Supplementary Table 5:**
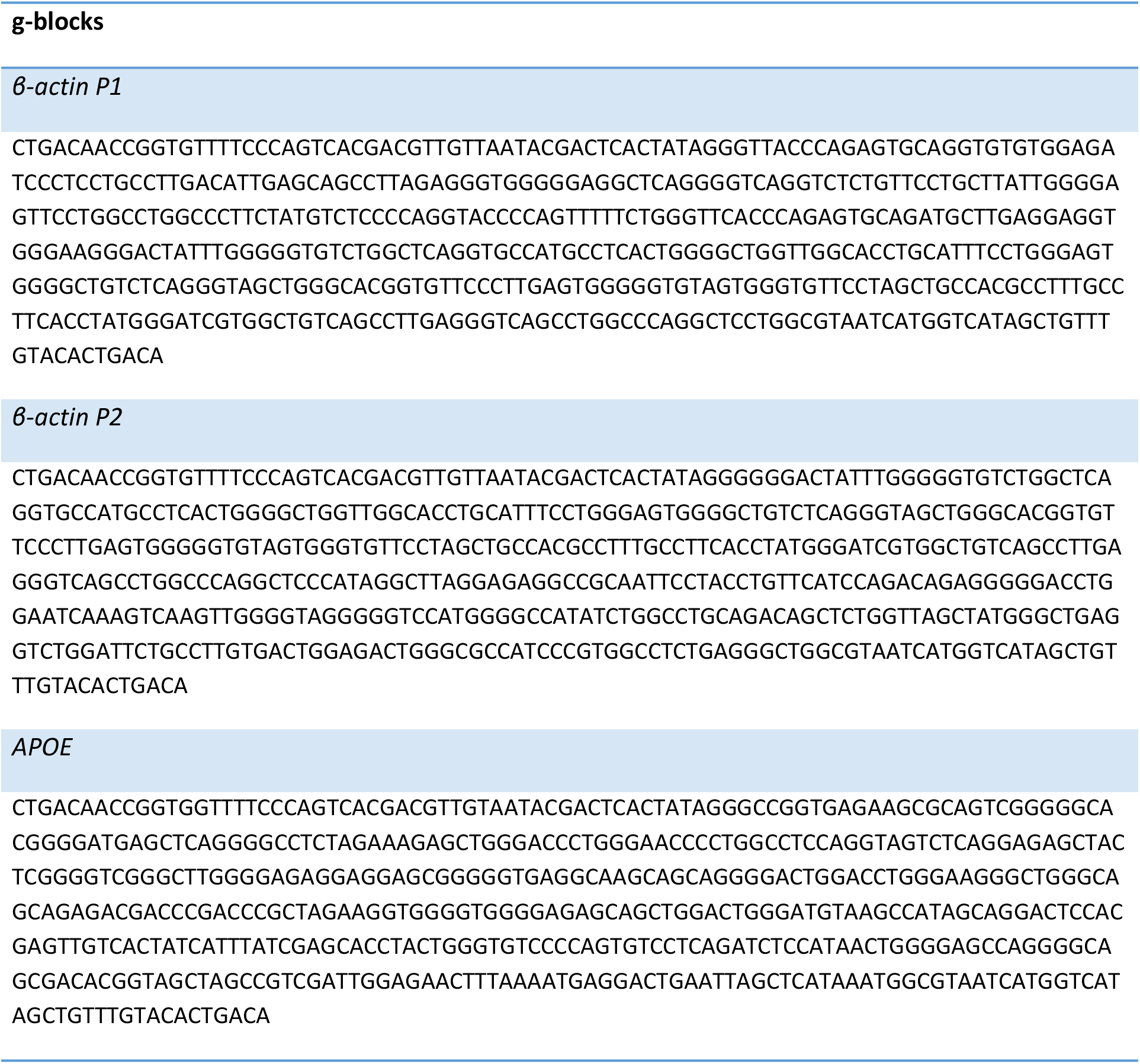
g-blocks sequences.

